# Comparative analysis of chemically and biologically synthesized iron oxide nanoparticles against Leishmania tropica

**DOI:** 10.1101/829408

**Authors:** Qurat ul Ain, Arshad Islam, Akhtar Nadhman, Masoom Yasinzai

## Abstract

The current study was carried out to compare the antileishmanial potential of the characterized chemically and plant mediated (*Trigonella foenum-graecum*) FeO-NPs (FeO-NPs) against *L.tropica* KWH23. The promastigotes and amastigotes mortality, ROS generation, and biocompatibility assessment were estimated in time-dose dependent manner by PDT. LED exposed bio-inspired FeO-NPs after 72 hours, express significant suppressive effects by exhibiting 50% inhibitory concentration (IC_50_) 0.001572±0.02μg/ml and 0.011408±0.02μg/ml for promastigotes and amastigotes, respectively. Leishmanial DNA damage and membrane integrity caused by FeO-NPs were owe to high production of reactive oxygen species (ROS). By applying different ROS scavengers (mannitol and sodium azide) hydroxyl radical and singlet oxygen were found as main moieties for cell death by giving quantum yield 0.15 and 0.28 through chemically and green synthesized FeO-NPs, respectively. The light exposed green synthesized nanoparticles were found biocompatible on human red blood cells (RBCs), by exhibiting LD_50_ value 2659μg/mL. From these findings, it can be concluded that plant mediated FeO-NPs can be used as promising antiprotozoal agent.

## Introduction

Leishmaniasis is a vector born protozoal disease caused by parasite belonging to genus *Leishmania*. It is worldwide infectious disease, causing increase rate of mortality and morbidity in different region mainly in tropics and subtropics areas (Alvar, Yactayo, & Bern, 2006). World Health Organization (WHO) reported it as one of serious most parasitic diseases and ranked second after malaria. The life cycle of parasitic protozoans is complex and characterized by three major developmental stages; promastigotes, metacyclic-promastigotes and amastigotes (Ullah et al., 2016).

*Leishmania* parasite have extraordinary ability to survive in alternative forms in mammalian macrophages and sand-fly guts. During leishmanial infection, the parasites are phagocytized by the macrophages of host individual as a result generation of reactive oxygen species (ROS) occurs. ROS like superoxide, hydrogen peroxide (H_2_O_2)_ and hydroxyl radicals have ability to produce oxidative stress by causing damage to macromolecules including proteins, lipids, and nucleic acids thus effecting the cell viability, integrity and making the cell wall fragile (Palmieri & Sblendorio, 2007). The *Leishmania* pathogens are very sensitive to these oxygen derived moieties as they are highly reactive and unstable. In order to cope with these reactive oxygen species, *Leishmania* produces enzymes such as acid phosphatase, peroxiredoxins, and lipophosphoglycan which act as scavenger of ROS release by macrophages (Allahverdiyev et al., 2013). This problem can be overwhelming by releasing more ROS extracellularly to the extent that it can cause the death of pathogens, while safer for human cells.

In endemic areas, currently practiced means of diagnosis, vector control, and chemotherapies have become outdated because of ineffectiveness. There are various complications associated with currently available modalities(Pentostam, Glcantime, pentamidine and amphotericine B) to control leishmaniasis (Yasinzai, Khan, Nadhman, & Shahnaz, 2013).These drugs are overwhelmed because of possible side effects, complicated administration and not cost effective as well (Shah, Khan, & Nadhman, 2014).

Taking into account these mentioned complications there is utmost need of modern era to explore natural products as an alternative synthetic antibiotic agents (Gleiter, 2009). By understanding the nature of parasite interaction with the mammalian host and the biology behind the cellular and molecular mechanism responsible for parasitism is expected to be helpful in rational development of more effective and efficient therapies.

There are several studies that demonstrate the bactericidal effects of FeO-NPs. Antibacterial activity of FeO-NPs against *Staphylococcus aureus, Salmonella enterica, Escherichia coli* and *Proteus mirabilis* have been reported (Naseem & Farrukh, 2015). It is known that production of reactive oxygen species (ROS) is main mechanism that is responsible for the antibacterial properties of FeO-NPs. Reactive oxygen species like superoxide radicals (O2), hydrogen peroxide(H_2_O_2_) and singlet oxygen cause oxidative stress that cause damage to macromolecules including proteins and nucleic acid in bacteria. In some cases, certain nanoparticles are activated by photo dynamic therapy (PDT), where ROS production is stimulated under the exposure of photon particles (light) (Matějka & Tokarský, 2014).

Chemical route of synthesis involves the use of highly toxic and hazardous chemical as the reducing and capping agents followed by generation of hazardous byproducts. Their applications in the field of nanomedicine is highly limited because it may cause some harmful effects. Hence, in the field of nanotechnology, there is critical need to introduce most reliable, effective, and eco-friendly approaches to synthesized nanoparticles. To fulfil this need, the use of natural products like biological systems to synthesize nanoparticles is good alternative. In green synthesis of nanoparticles plants phytochemicals can be used as reducing and capping agents, it refers as more effective and less harmful approach.

Antibacterial, antifungal, antiviral, and anti-carcinogenic efficacy of FeO-NPs have been observed and reported but to date, there is no reported study that demonstrate their anti-leishmanial efficiency on *Leishmania* parasite. It is the preliminary study that investigate the antileishmanial effects of FeO-NPs under different conditions (dark and LED exposures).

## Material and methods

### 2.1. Chemicals and Reagents

Iron chloride hexahydrate (FeCl_3_.6H_2_O), sodium hydroxide (NaOH), sodium bicarbonate, fetal bovine serum (FBS), HEPES, ethylenediaminetetraacetic acid (EDTA), sucrose, iron chloride tetrahydrate (FeCl_2._4H_2_O), sodium dodecyl sulphate, Tris-HCL, triton X-100, proteinase K solution, phenol, chloroform, diphenylisobenzofuran (DPBF) were purchased from Sigma-Aldrich^®^(St. Louis, MO, USA). Deionized water (MilliQ^®^) was used throughout the experiments. All the solvents and regents were at of analytical grades and used as received.

### 2.2. Synthesis of FeO-NPs

FeO-NPs were synthesized either through chemical or biological synthetic procedure, in order to assess and compare their antileishmanial potential.

#### 2.2.1. Chemical synthesis

Stock aqueous solutions of 0.2M FeCl_3_.6H_2_O) and 0.1M FeCl_2_.4H_2_O metal salts was prepared as iron source by dissolving respective chemicals in 100m distilled water. It was heated at 70 °C for 30-35 min in water bath, followed by the drop wise addition of 10 mL of 1 M NaOH _(aq)_ solution. An instant formation of black precipitate was observed but the stirring was allowed for 1-2 h. The black precipitates were separated by centrifugation (WiseSpin, CF-10) at 10,000 rpm for 10 min and washed 4-5 times with distilled water to filter-off the impurities. The FeO-NPs were harvested by drying the precipitates in oven (Oven Daihan Labtech LDO-060E) at 60 °C for overnight and subjected to calcination at 400°C for 3 h.

#### 2.2.2 Green synthesis

Stock aqueous solutions of 0.2M FeCl_3_.6H_2_O) and 0.1M FeCl_2_.4H_2_O metal salts was prepared as iron source by dissolving respective chemicals in 100m distilled water<u>. Bio solvent was extracted from</u>*<u>Trigonella</u> <u>foenum-graecum</u>* <u>(fenugreek) plant. For the preparation of extract, 10g of seeds were rinsed with distilled water to remove the surface impurities and then boiled in 100ml of water at 70°C for 10 minutes. The extract was filtered by using whatman filter paper (pore size is 90 nm) and centrifuged at 10,000 rpm for 10 minutes to obtain the supernatant. The bio solvent was added to iron precursor solution in 1:2 volume, and kept at 70°C with constant stirring for 30 minutes. At the completion of reaction, the dark black precipitate were formed at pH 11. Precipitates were washed with distilled water for two times to remove all the ions by centrifugation</u>(WiseSpin, CF-10) <u>at 10,000 rpm for 10 min. The synthesized bio-inspired nanoparticles were dried in oven</u> (S.R Traders, DHG-9053A) <u>at 80°C for overnight.</u>

### 2.4. Characterization of FeO-NPs

The synthesized nanoparticles were characterized by various techniques. The optical properties of nanostructure were studied through UV-Vis spectrophotometer (Shimadzu UV-1601 spectrophotometer). The spectra was recorded between the ranges of 200-500 nm. The various reducing phytoconstituents and chemical composition of FeO-NPs was analyzed by Fourier Transform Infrared (FT-IR) spectroscopy (IRTracer-100 FT-IR Spectrophotometer) in the range of 3000–300 cm^-1^, using anhydrous potassium bromide (KBr) pellet. The grain size and phase variety of nanostructure (FeO-NPs) were obtained by X-ray diffraction analysis (Shimadzu, XRD-6000).

### 2.5. *Leishmania* cell culture

The *L.tropica* KWH23 promastigotes were provided by Department of zoology, university of Peshawar, Pakistan. Parasites were cultured in M199 medium supplemented with 10% fetal bovine serum x (FBS Sigma, St Louis, MO), sodium bicarbonate (Sigma, USA), streptomycin (Bio Basic Inc.) and HEPES buffer (PAA laboratories Gmbh).

### 2.6. Nanoparticles solution

The stock solution of FeO-NPs (green and chemical) was obtained by dispersing nanoparticles (1mg) in ultrapure water (1 ml). To avoid the agglomeration, vertexing was done for 1-2 minutes, followed by sonication for 10 minutes to form homogeneous suspensions.

### 2.7. Determination of time dependent effects of FeO-NPs on proliferation of promastigotes

In vitro, anti-promastigotes assay was assessed by using promastigote culture of *Lesihmania tropica* by applying individual concentrations of green and chemically synthesized FeO-NPs in 96 well culture plate. Promastigotes in the exponential phase were seeded at 1 × 10^8^ /100 *μ*L cells per well of microtiter plate. Afterwards, the parasites were treated with serially diluted nanoparticles (10μg/ml, 5μg/ml, 2.5 μg/ml, 1.25 μg/ml, 0.625 μg/ml, 0.3125 μg/ml, 0.15625 μg/ml, 0.078125 μg/ml). The experiments was divided into two assays, first was exposed to LED light for 10 minutes, and while second was without light exposure then both groups were incubated for 72 hours in darkness at 25°C. The leishmanial culture without any treatment (light and nanoparticles) was taken as control. Antileishmanial activity was evaluated by using Neubauer chamber (Micros, Carinthia, Austria) under inverted microscope (Micros, MC700 - Austria), counting the viable cells at the incubation time period of 24, 48, and 72 hours.

### 2.8. Anti-leishmanial activity against amastigotes

FeO-NPs (chemical and green) were further evaluated for leishmanicidal activities against amastigotes. In 96 well plate medium M199 containing culture of amastigotes (2×10^6^ cells/mL; 2 ml/well) was added in each well, afterward supplemented with serially diluted FeO-NPs (chemical and green). Similar to promastigotes, two conditions were applied by dividing the experiment into two groups, first was exposed to LED light for 10 minutes while second was proceeded under dark conditions. Following incubation for 72 hours, antileishmanial potential of all the analyzed concentrations was determined by counting the viable amastigotes by Neubauer chamber after staining with trypan blue.

### 2.9. Cytotoxicity assay on human erythrocytes

Hemolytic activity was evaluated by taking the blood (A^+^) of an healthy individual (Human red blood cell’s approved by SA-CIRBS, International Islamic University, Islamabad, Pakistan review committee on November 17, 2016). The blood was subjected to centrifugation at 3000 rpm for 3 minutes, the pellet containing erythrocytes was further washed with PBS. Then PBS diluted erythrocyte suspension was treat with nanoparticle. Experiment was carried out under two conditions. One group was exposed to LED light for 10 minutes while second was under dark conditions and incubated the both groups at 37°C for 3 hours. Red blood cells, treated with 0.1% triton X-100 was taken as positive control while erythrocytes suspended in phosphate buffer, without nanoparticles exposure was taken as negative control. After treatment, erythrocytes suspension was centrifuged at 6000rpm for 10 minutes and cell lysis was estimated spectrophotometrically (576nm). The evaluations were obtained by the percentage of hemolysis compared with the results obtained from the positive and negative controls. 

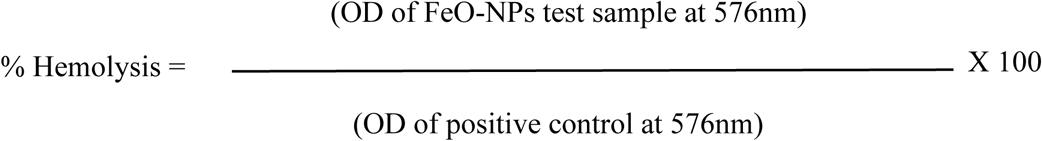

### 2.10. Reactive Oxygen Species (ROS) Quantification

The *in vitro* reactive oxygen species (ROS) generated by FeO-NPs was calculated by chemical oxidation of 1, 3-diphenylisobenzofuran (**DPBF**) in ethanol solution. Briefly, 20μg/ml of FeO-NPs were treated with 1.5ml of 0.15 mM solution of DPBF in a sealed quartz cuvette, followed by LED light exposure. The oxidation and photo bleaching effects of DPBF was estimated by measuring the absorbance by UV-Vis Spectrophotometer (Shimadzu UV-1601 spectrophotometer) at 410nm at the time interval of every 30 seconds up to time period of 350 seconds. For measuring the decaying rate of photosensiting process, the graph was plotted between natural logarithm of absorbance against the irradiation time. Methylene blue was used as a standard. To evaluate main ROS moieties, ROS scavengers like 0.1mM sodium azide and 0.1 mM mannitol were used.

### 2.11. Qualitative and quantitative analysis of FeO-NPs against *Leishmania* DNA

DNA extraction from *Leishmania* (nanoparticles treated and untreated) was performed by adding 100µl lysis buffer (2 mM ethylenediaminetetraacetic acid (EDTA) (pH 8.0), 0.3 sucrose, 10 Mm sodium dodecyl sulphate, 10 mM Tris-HCL (pH 7.5), 1% triton X-100, 8.5 µl proteinase K solution), afterward incubated at 60°C for 2 h in waterbath. After incubation, equal ratio of phenol: chloroform (1:1) was added to the above reaction, followed by centrifugation at 10,000 rpm for 10 minutes. Supernatant, the aqueous layer containing the DNA was separated, 1ml of chilled isopropanol was added. The resulted solution was again centrifuged at 10,000 rpm for 10 min. The pellet containing the DNA was washed with 70% ethanol and centrifuged for 2 min. The precipitated DNA was suspended in 50 µl of Tris-EDTA buffer (0.5 M EDTA, 1 M -HCL). For qualitative analysis, reaction was performed in two groups. In first group, *Leishmania* culture was treated with FeO-NPs under light and dark conditions, while in second group, isolated *Leishmania* DNA was treated with nanoparticles under both conditions.

For qualitative analysis DNA was extracted from the treated *Leishmania* (LED light and dark) by the above method. All the DNA samples including treated *Leishmania* DNA and extracted *Leishmania* DNA (treated with nanoparticles) were electrophoresed by using 1% agarose gel at 100V. For DNA visualization, 0.7µg/ml of ethidium bromide was used. Quantitative analysis was done by nanodrop: the nanodrop determine the quantity and purity of the DNA based on the absorbance at A260.

### 2.12. Apoptosis, necrosis, and membrane permeability evaluation

To quantify the cell death, leishmanial cells were treated with FeO-NPs in two groups, first group was exposed to LED light for 10 minutes at 25°C, while second group without any light exposure, both assays were incubated for overnight under dark conditions. After incubation time period, phosphate buffer solution (PBS) was used for washing of treated groups, followed by centrifugation at 6000 rpm for 10 minutes. Pellet was suspended again in the PBS and then divide the treated groups further into two groups for the staining with fluorescent dyes (ethidium bromide and acridine orange). One groups was stained with 100µg/mL acridine orange while second group was stained with both AO and EB in in 3:1 (Nadhman et al., 2014). Florescent microscope was used to measure the fluorescence.

To assess the membrane permeability SYTOX green dye was used. FeO-NPs treatment was same just like for apoptotic and necrotic evaluation, while washing was done with HBSS. The nanoparticles treated cells were stained with 1μM SYTOX dye for 15 minutes under dark condition. Then fluorescent intensity was measured through fluorescent microscope. Triton X-100 (0.1%) was used as a control to cause complete membrane integrity.

### 2.13. Analysis of data and statistics

In recent study, all the experiments were performed in triplicates and findings are presented as mean ± standard deviation. Significant differences assessment was done by the Student’s t-test through SPSS software.

## 3. Results

### 3.1. Characterization of FeO-NPs

#### 3.1.1 UV-Vis spectrophotometer

Synthesis of FeO-NPs (green and chemical routes) was physically observed by color change from yellow to black and confirmed by UV-Vis spectrophotometer. This color change was because of Surface Plasmon Resonance phenomena. This technique specifies the rate of metal reduction as well as progress of nanoparticle synthesis. Fig.3.1 shows the absorbance spectra of FeO-NPs synthesized from chemical and green route. Characterized surface Plasmon resonance for green and chemically synthesized FeO-NPs shows sharp spectral band at 267 nm and 268 nm, respectively.

**Fig.3.1.**
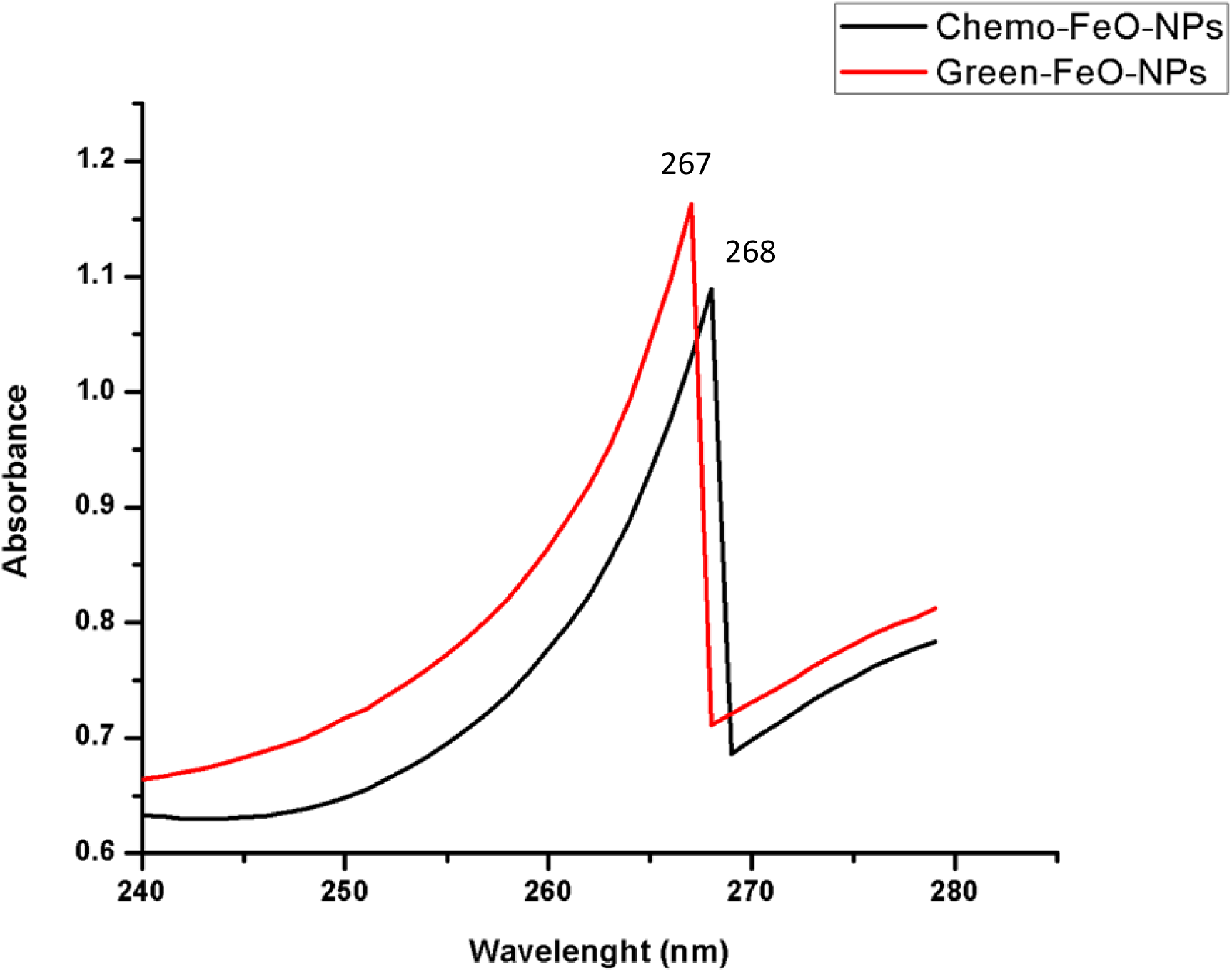
UV spectra of synthesized FeO-NPs

#### 3.1.2. FT-IR analysis of FeO-NPs

Fig.3.2 (a) show FTIR spectrum of green synthesized FeO-NPs and *Trigonella foenum-graecum* seeds extract. Spectrophotometer (IRTracer-100 FT-IR) in the range of 300 cm-1 to 3000 cm-1 was used to observe the infrared absorption peaks indicating the various functional groups acting as reducing and capping agents.

**Fig. 3.2 (a).**
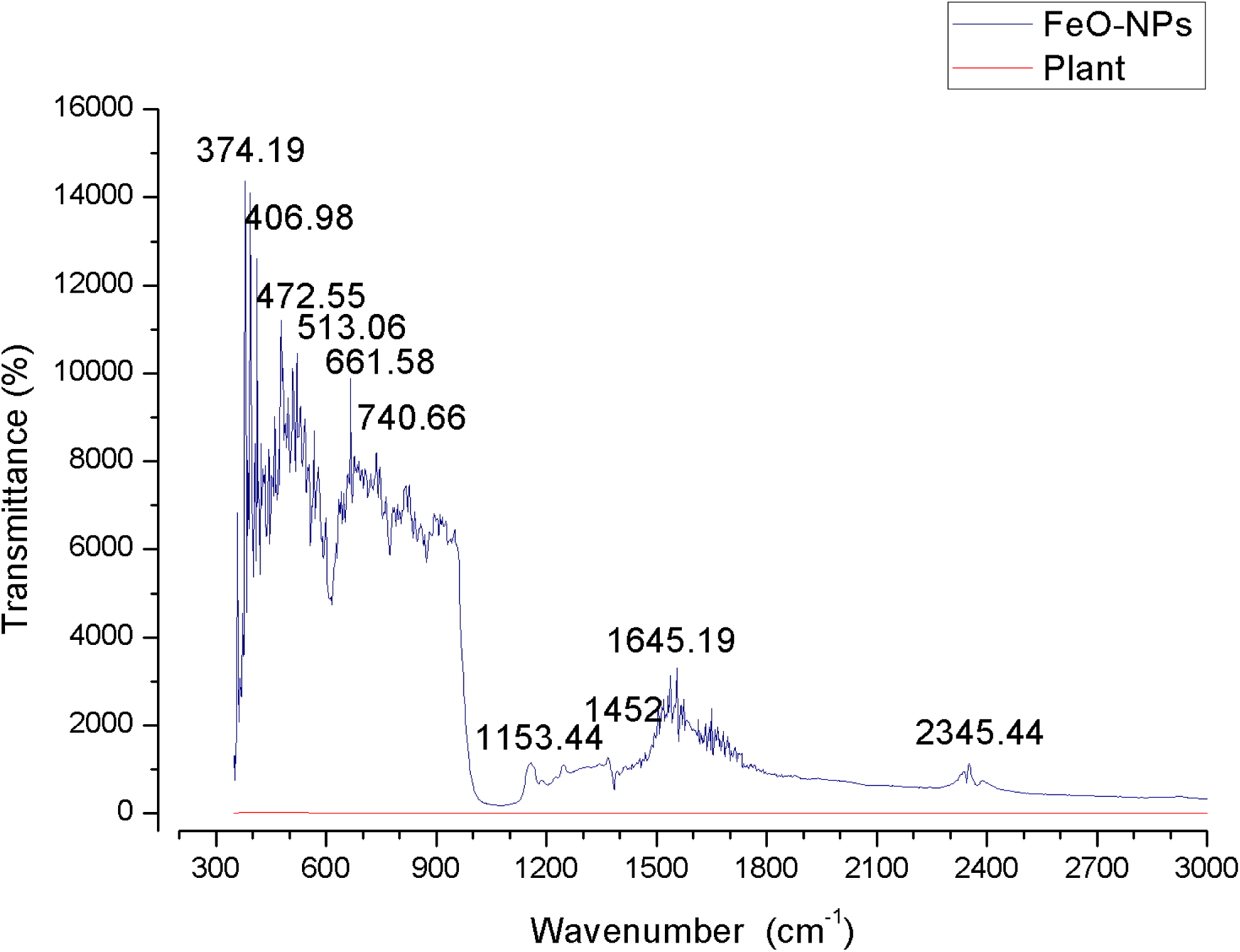
FTIR Analysis of Green synthesized FeO-NPs

The FT-IR spectra of green synthesized FeO-NPs manifest prominent peaks at 374.19 cm^-1^, 406.98 cm^-1^, 472.55 cm^-1^, 513.06 cm^-1^, 661.58 cm^-1^, 740.66 cm^-1^, 1153.44 cm^-1^, 1452.39 cm^-1^, 1645.19 cm^-1^, 2345 cm^-1^ (Fig.3.2 a)

The IR peaks at the less wave number (<700cm-1) are responsible for the vibrations of Fe-O bonds of iron oxide (Cornell & Schwertmann, 2003). The intense absorption band at 513.06 cm-1 attributed to the Fe-O stretching, which relates to the magnetite phase (Mahdavi et al., 2013) and also the peak at 661.58 cm^-1^ indicates the Fe-O vibrations of FeO-NPs, thus confirming the synthesis of magnetic FeO-NPs. The peak at 1452.39 correspond to C-C groups of aromatic rings that are present in plant extract and also the absorption band at 1648.19 cm-1 is associated with stretching vibration of carbonyl groups. The peaks at 2345 show the presence of N-H bindings of amide group.

From FTIR analysis, soluble components present in the plant extract could have considered as reducing and stabilizing agents, preventing the nanoparticle aggregation in solution form, and thus playing vital role in synthesis of FeO-NPs.

Fig. 3.2 (b) shows the spectra of chemically synthesized FeO-NPs. The prominent absorption bands were observed at 551cm^-1^ 619.14 cm^-1^, 775.86 cm^-1^, 1107.41 cm^-1^, 1382.96 cm^-1^, 1641.42 cm^-1^, and 2069.28 cm^-1^. The bands at 619.14 cm^-1^, 775.86 cm^-1^, 1107.14 cm^-1^ correspond to O-H stretching vibration, carboxyl group, alkenyl C=C stretch respectively. The characteristic IR vibrations of FeO-NPs at 551 cm-1 relates to the bending and stretching vibrations of Fe-O bond, which confirms the synthesis of FeO nanoparticles.

**Fig. 3.2 (b).**
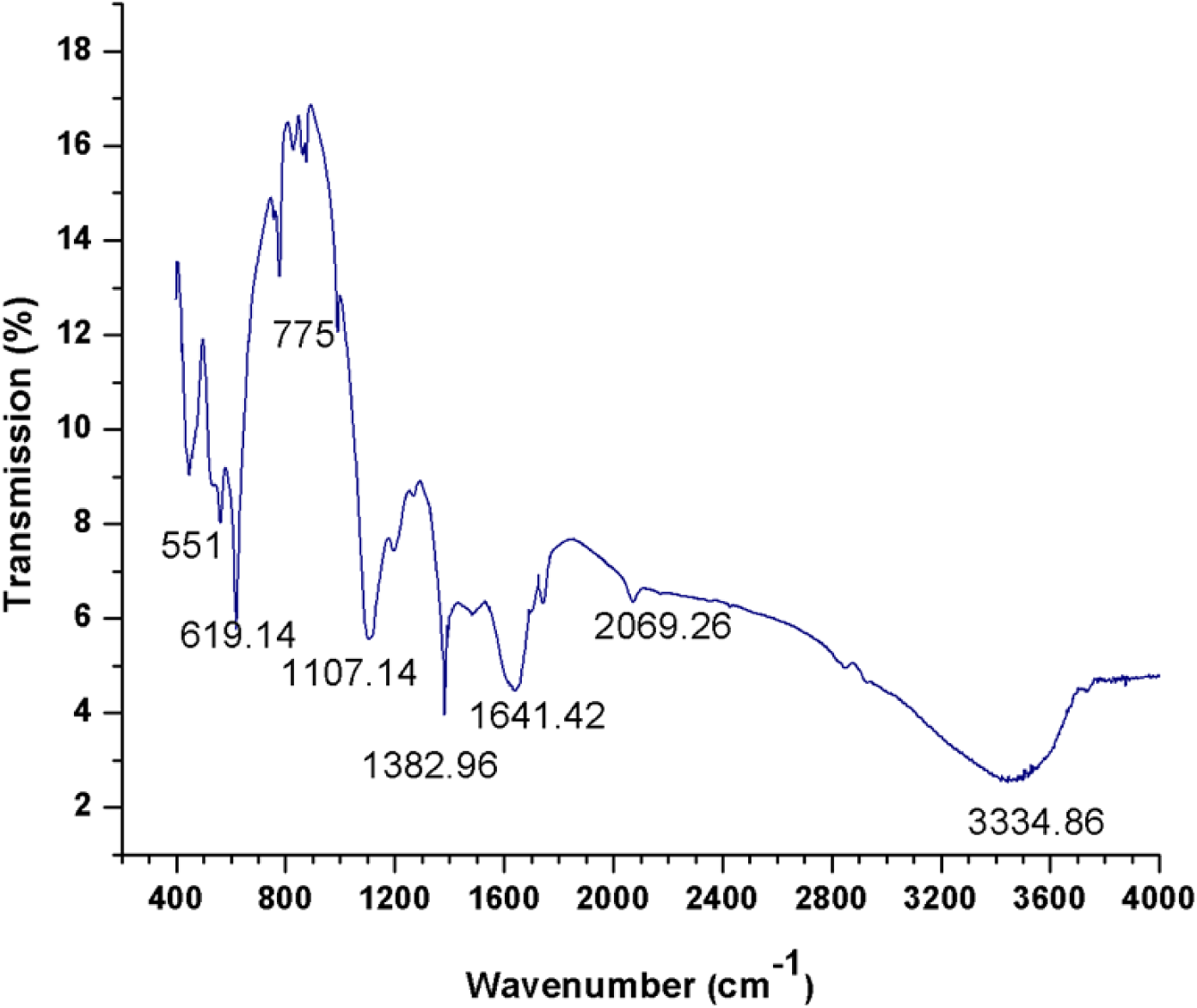
FTIR Analysis of Chemically synthesized FeO-NPs

### 3.2. Dose- time dependent in-vitro anti-promastigotes activity

For evaluation of antileishmanial efficacy, the viability of promastigotes was checked in both control and test groups at various time intervals (24h, 48h, and 72h). In figure 3.4. Significant decrease in viability was seen in the treated groups as compare to control. This effect was examined during all analyzed incubation time periods. Beside this determination, there was a considerable decrease in promastigotes survival was observed in LED exposed, NPs (chemical and green) treated groups as compare to without LED exposed NP’ s treated groups. Fig.3.4 and Fig.3.5.clearly shows, the effects of FeO-NPs on the viability of promastigotes, under the LED light were more significant than they were in the dark conditions.

**Figure 3.4.**
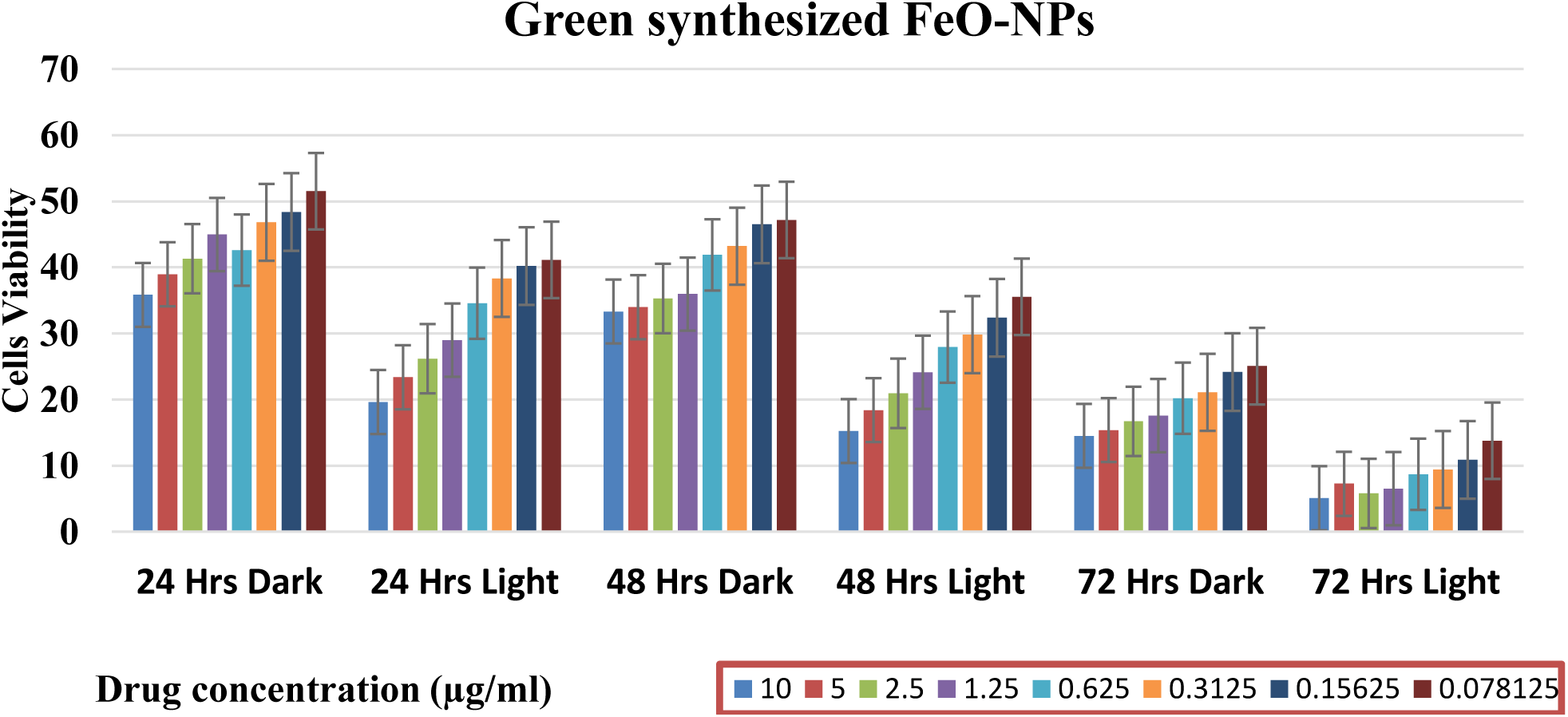
Anti-leishmanial activity against promastigotes of *Leishmania tropica* of green synthesized FeO nanoparticles

**Figure 3.5.**
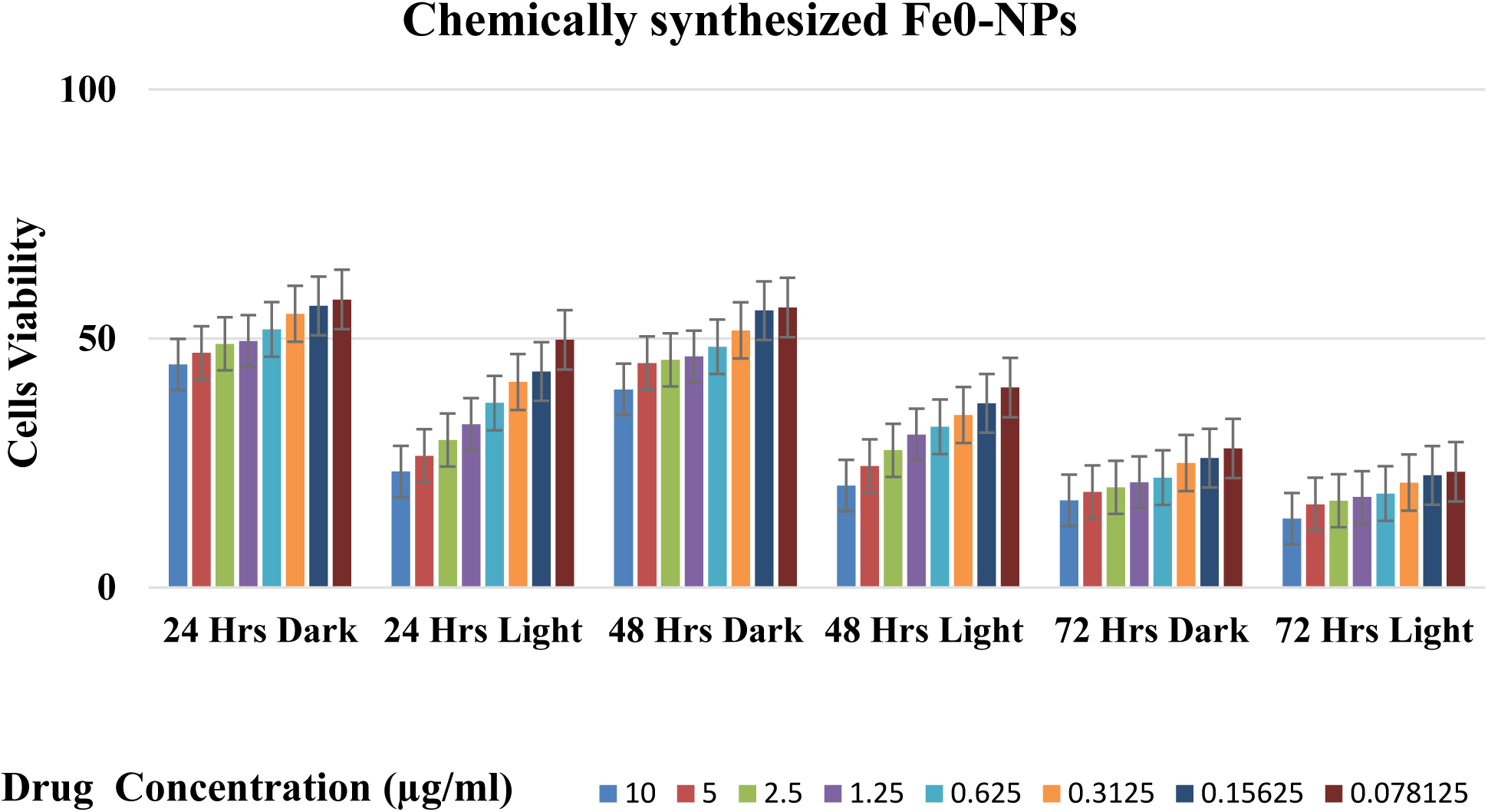
Anti-leishmanial activity against promastigotes of *Leishmania tropica* of chemically synthesized FeO nanoparticles

Beside these determinations, there was significant increase in mortality rate was observed in green synthesized FeO-NPs treated groups. This assessment was further confirmed by estimating the ROS production. Reactive oxygen species are highly reactive that can cause oxidative stress by causing biochemical and morphological changes. The maximum mortalities were observed in LED exposed, green synthesized nanoparticle treated group, after 72h of incubation at highest dose concentration with IC_50_ value of 0.001572±0.022 µg/ml. Further IC_50_ values after 24 and 48h was 0.020274±0.015 µg/ml and 0.018577±0.016 µg/ml, respectively. While under dark condition, nanoparticles (green) treated groups show IC_50_ values 0.022512±.014 µg/ml, 0.02070±.014 µg/ml, and 0.001512±0.017 µg/ml at incubation period of 24h, 48h, and 72h respectively.IC_50_ values for chemo-nanoparticles treated groups, under dark conditions are, 0.025385±.014, 0.022522±.014, 0.001938 ±0.016 at incubation period of 24h, 48h, 72h respectively. While for LED exposed groups, IC_50_ values are 0.022705 ±.015, 0.018324 ±.015, 0.001938 ±0.017, respectively. (Fig.3.6.)

**Fig.3.6.**
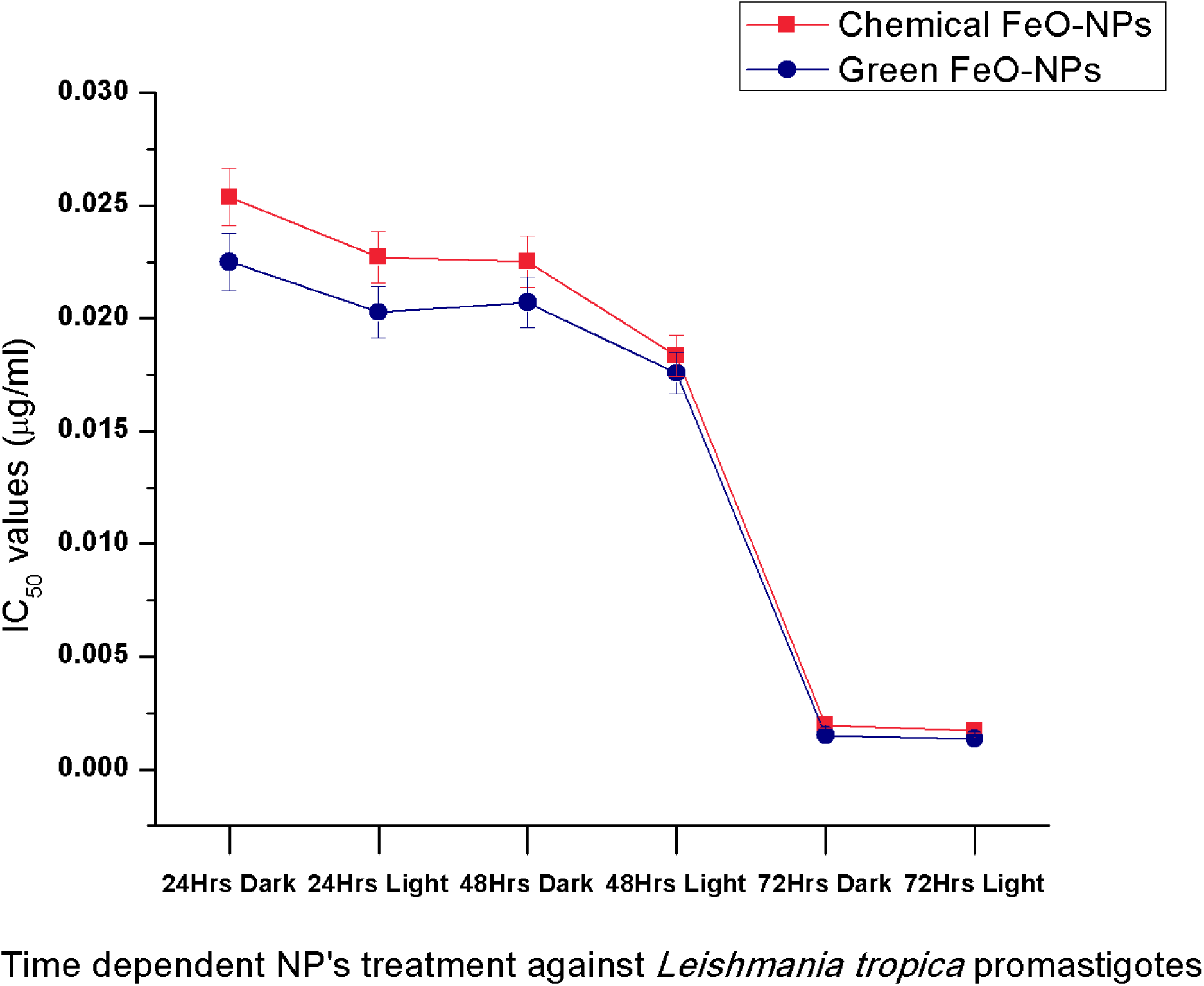
Statistical comparison of IC_50_ of Green and Chemically synthesized Fe-NPs against *Leishmania tropica* promastigotes

Thus these findings clearly conclude that FeO-NPs at lower dose can significantly kill the *Leishmania* cells and it may serve as favorable candidate for the treatment of fatal leishmaniasis.

### 3.3. Anti-leishmanial activity against amastigotes of *Leishmania tropica*

Leishmanial amastigotes were incubated with FeO-NPs (chemical and green) at differnet dilutions for 72hrs. Similar to promastigotes, the reaction was performed in two groups, one was exposed to LED light for 10 minutes while other was proceeded under dark conditions. Reduced survival of amastigotes was observed in both LED exposed and non LED exposed treated groups. Significant killing of cells was observed in light exposed, NPs treated groups. The green synthesized nanoparticles cause increased generation of ROS, as a result higher killing ability of amastigotes was observed,as compare to chemo-synthesized nanoparticles.

### 3.4. Biocompatibility of iron nanoparticles

FeO-NPs are promising candidate possessing broad spectrum antimicrobial properties. However, it is necessary to evaluate the biocompatibility of these nanostructures on normal cells to ensure their safer use. The FeO-NPs (chemical and green) were evaluated for their biocompatibility against human macrophages in dose dependent manners under dark and light (LED) exposure. The result clearly indicated that green synthesized nanoparticles in light exposure show biocompatibility at low concentrations, by exhibiting LD_50_ value 2659μg/ml. The experiment under dark conditions indicated that current photo-catalytic nanoparticles can be toxic at higher concentration. (Fig.3.10. Lethal dose concentrations (LD)

**Fig.3.7.**
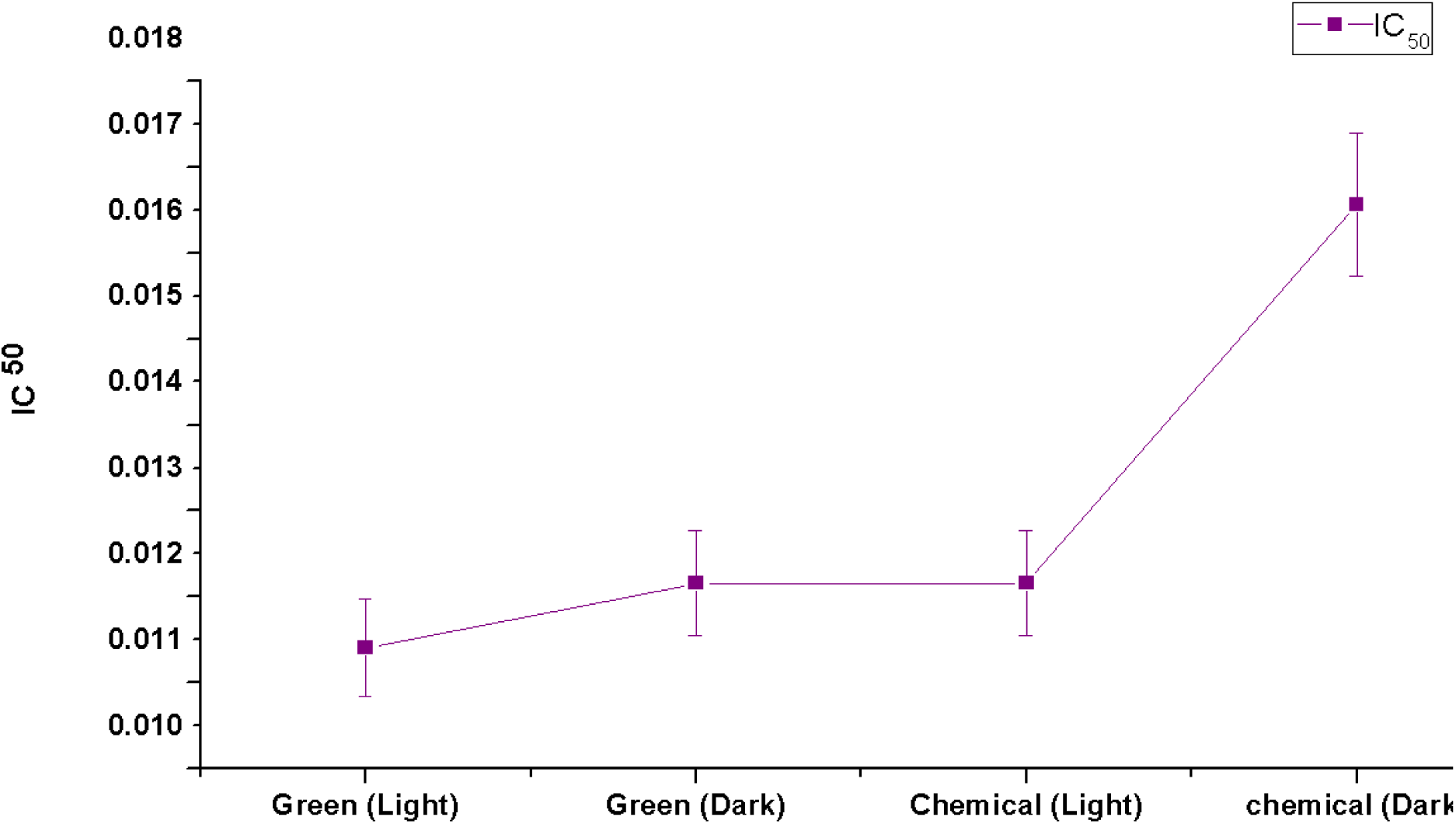
Statistical comparison of IC^50^ of Green and Chemically synthesized Fe-NPs against *Leishmania tropica* amastigotes.

**Fig.3.8.**
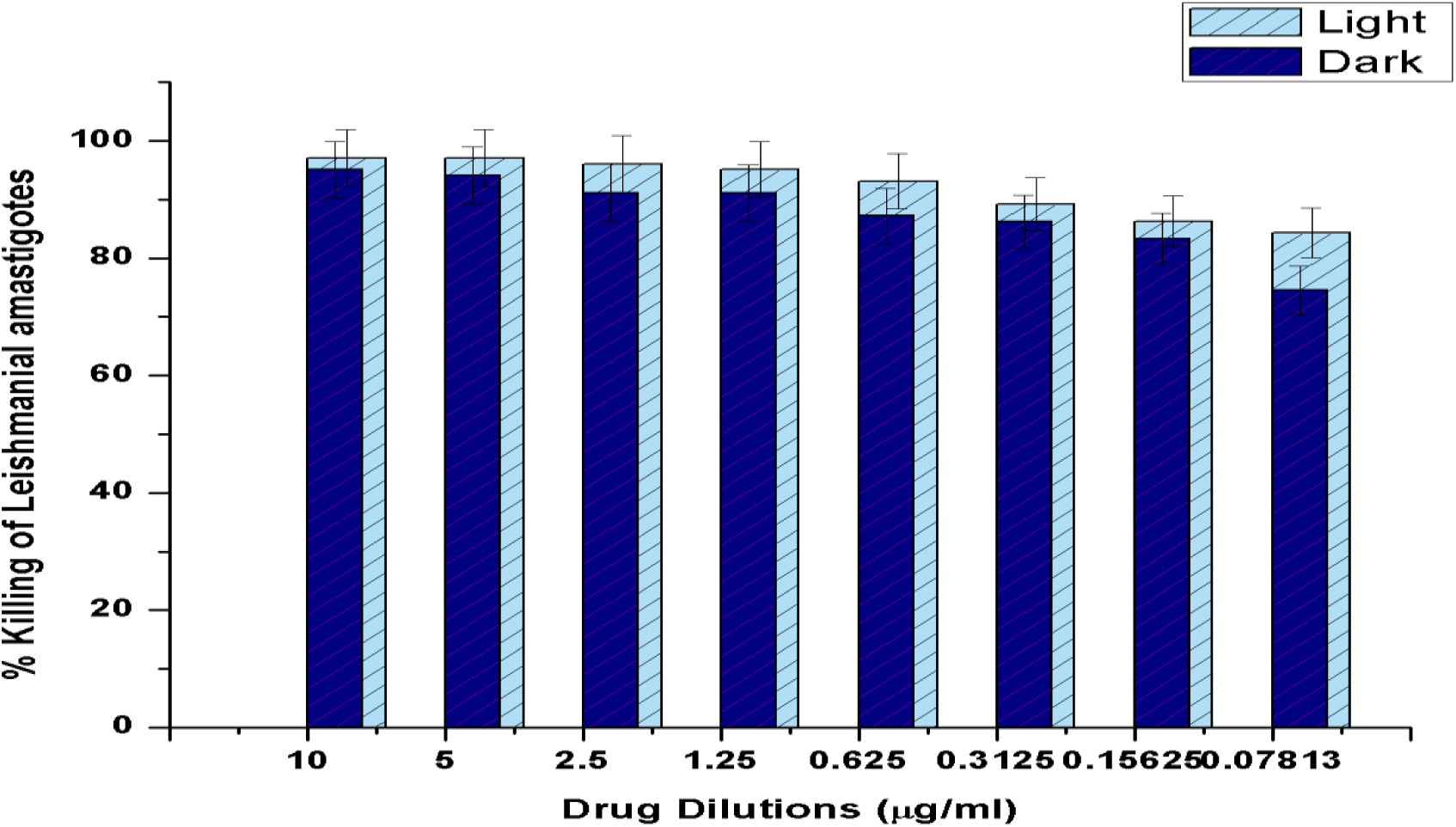
Activity of FeO-NPs (Green) against amastigotes of *Leishmania tropica*

**Fig. 3.9.**
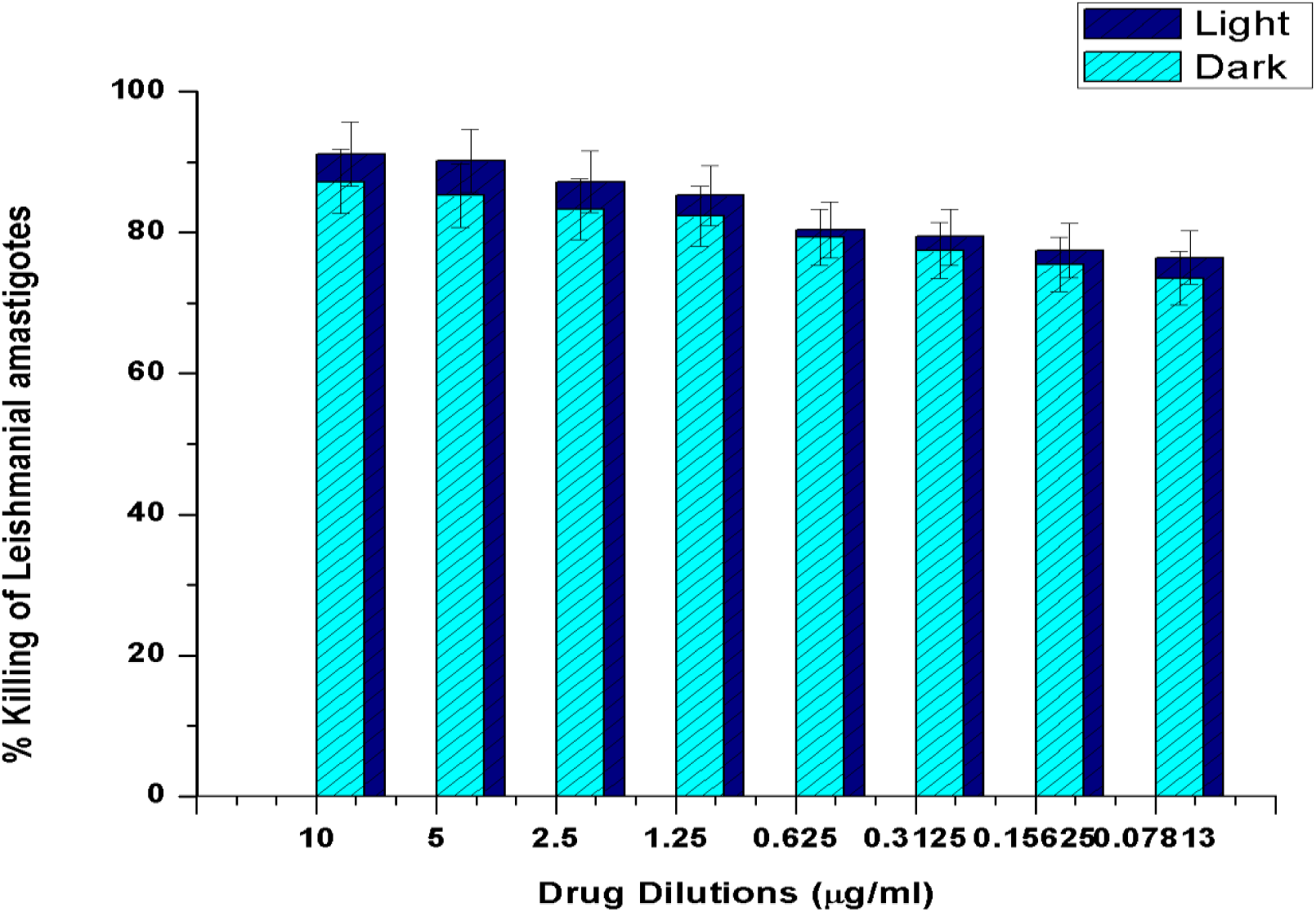
Activity of FeO-NPs (chemical) against amastigotes of *Leishmania tropica*

**Fig.3.10.**
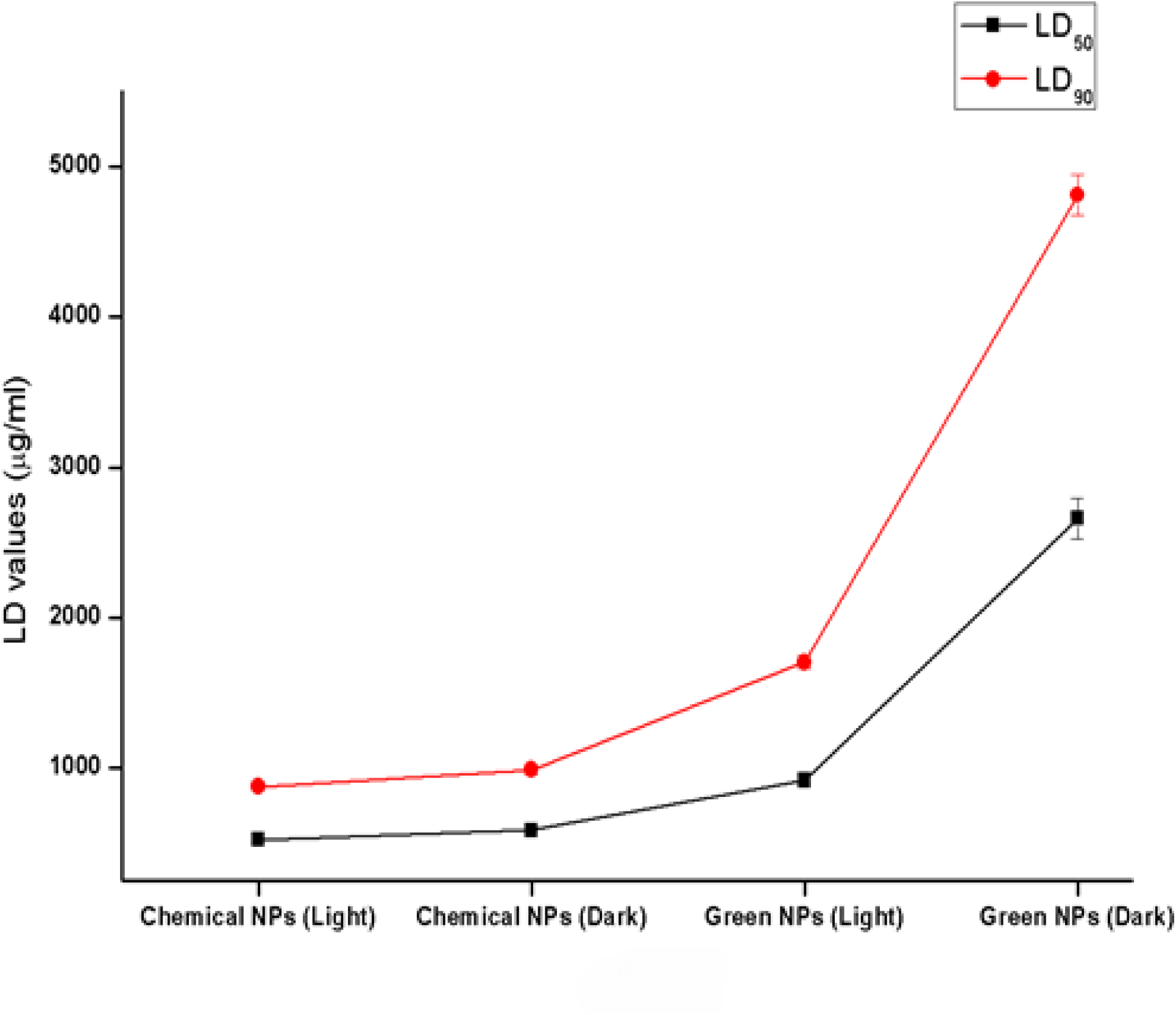
Lethal dose concentrations (LD) of Fe-NPs (green and chemically synthesized) against macrophages.

### 3.5. ROS quantification

Quantification of ROS produced by FeO-NPs was assessed by using 1, 3-Diphenylisobenzofuran (DPBF). It is a fluorescent probe that is used as indicator against some reactive oxygen species (hydrogen peroxide, singlet oxygen, hydroxyl radical and superoxide anion) by reacting in highly specific manner. Upon the action of reactive oxygen species, it readily transformed into 1, 2 dibenzoylbenzene (DBB)(Zamojc et al., 2017). By the action of ROS induced by photo-excitation of FeO-NPs, with DPBF, there was gradual decrease in its fluorescence. This chemical transformation was estimated by recording decrease in intensity of fluorescence after every 30 seconds through spectrophotometer at 410 nm.(Vankayala, Lin, Kalluru, Chiang, & Hwang, 2014). Comparative method was used for the estimation of quantum yield for ROS quantification in comparison with standard photosensitizer (methylene blue)(Xiao, Gu, Howell, & Sailor, 2011). 

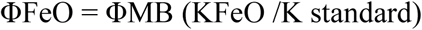

FMB is the quantum yield of methylene blue while FFeO is the quantum yield of photo-excited FeO-NPs. K_FeO_ and K _standard_ are the rate constant for the degradation of DPBF by photo-excited FeO-NPs and by standard photosensitizer, respectively. By assessing ROS generating ability of green and chemically synthesized nanoparticles, the quantum yield was estimated 0.28 and 0.15, respectively as compare to standard methylene blue (0.52).

Various ROS scavengers like mannitol and sodium azide were used for the evaluation of hydroxyl ions and singlet oxygen as the main ROS moieties responsible for causing leishmanial cells death.

### 3.6. Qualitative and quantitative analysis of *Leishmania tropica* DNA

DNA is an important cellular target of reactive oxygen species (ROS). The critical mechanisms behind the oxidative DNA damage involves Protein-DNA crosslinks, sugar and base lesions and double-strand breaks (Fu, Xia, Hwang, Ray, & Yu, 2014).

From ROS quantification it was found that both chemically and green synthesized nanoparticles have the ability to produce good quantum yield of ROS i.e. 0.15 and 0.26, respectively. Thus photoactive FeO-NPs have the ability to damage the *Leishmania* DNA in the presence of light. Figure 3.13 (a) show there is no DNA degradation under the dark conditions. Figure 3.13 (b) clearly indicate that upon light exposure FeO-NPs (green and chemical) have ability to degrade DNA by producing reactive oxygen species (ROS).

**Fig.3.11.**
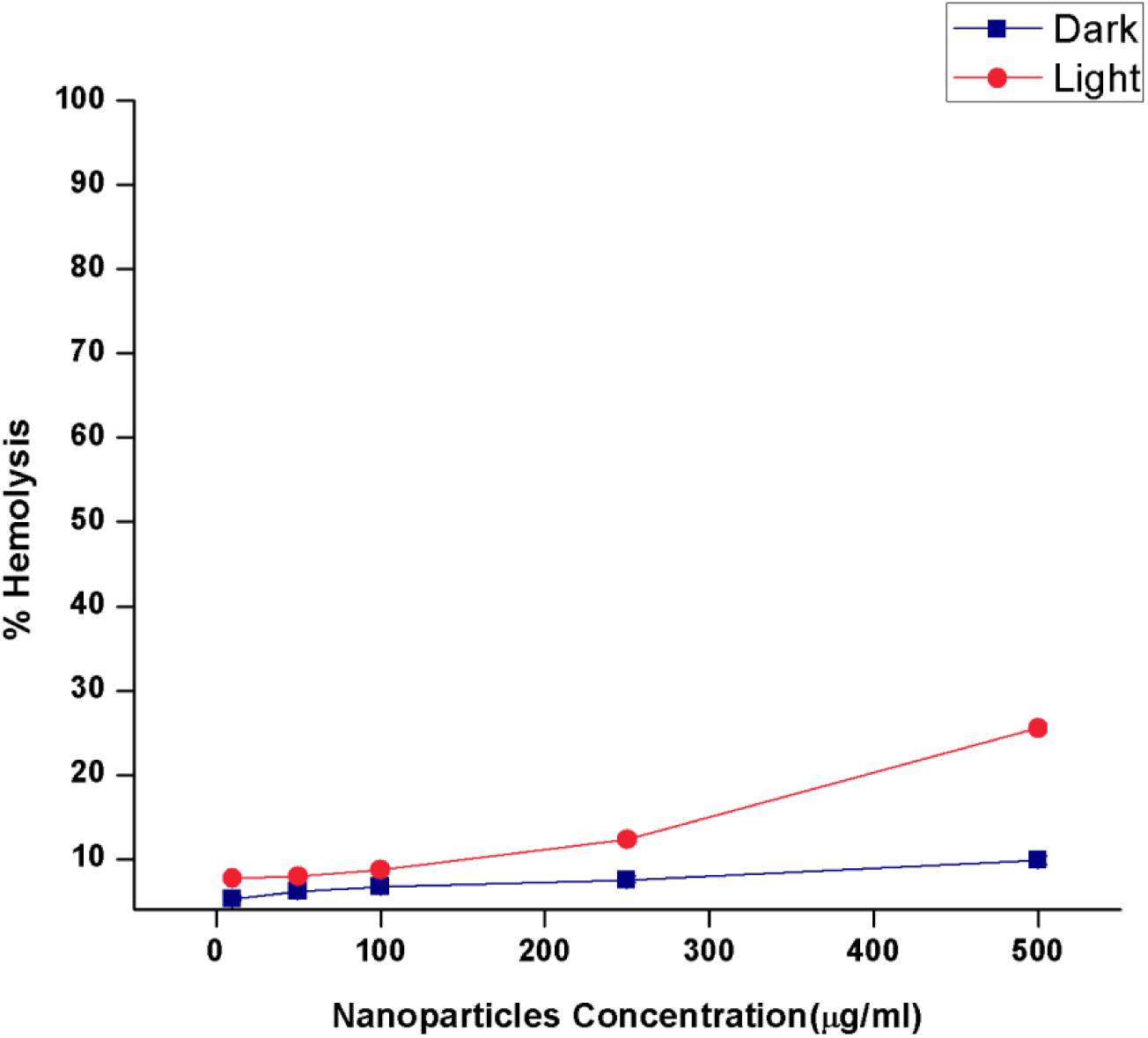
Biocompatibility of Green synthesized FeO-NPs against human blood

**Fig.3.12.**
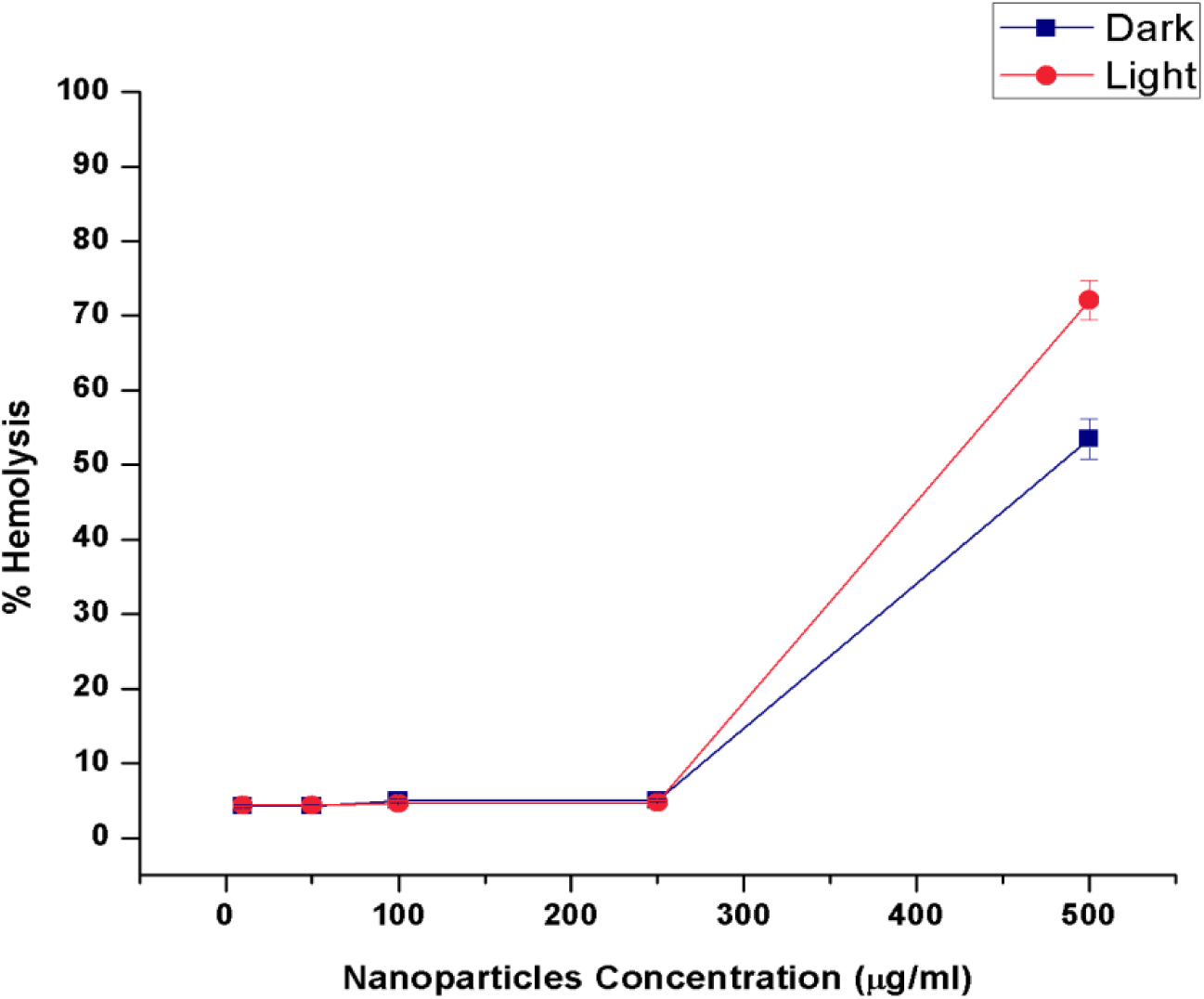
Biocompatibility of chemically synthesized FeO-NPs against human blood

**Fig. 3.13 (a):**
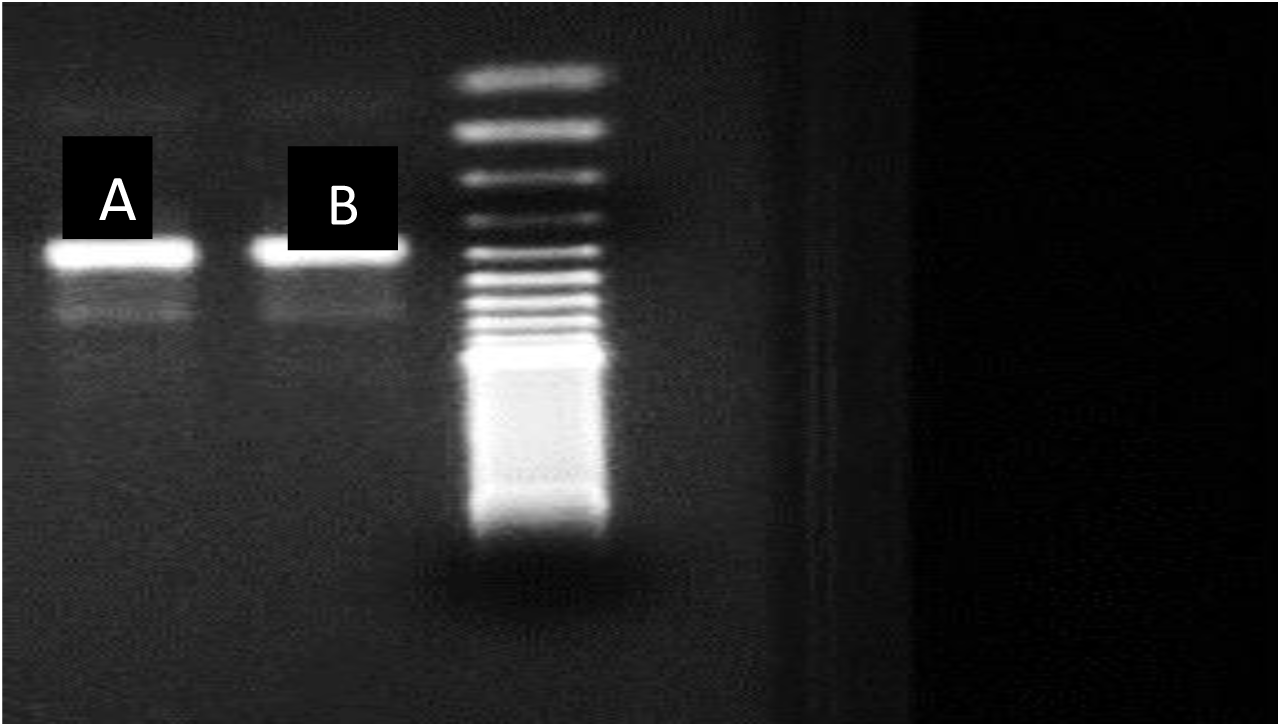
Gel electrophoresis of Leishmania DNA. (A) Control (untreated) Leishmania DNA; (B) treated Leishmania under dark conditions and its DNA.

**Fig. 3.13 (b):**
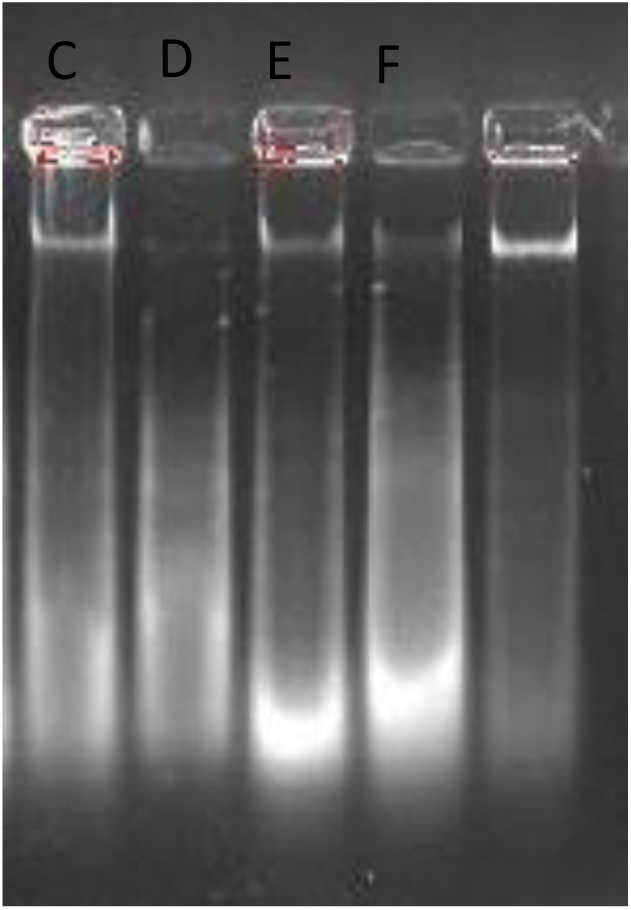
Notes: (C) (D) DNA extracted from treated Leishmania cells with FeO-NPs (green and chemical), on light exposure; (E) (F) DNA extracted from Leishmania cells and then treatment with FeO-NPs (green and chemical) and exposed to light.

For quantitative analysis, the extracted *Leishmania* DNA, without any exposure to nanoparticles was quantified as 618.3 ng/μl through nanodrop spectrophotometer (Nanodrop ND-1000 Spectrophotometer). Then this extracted DNA was treated with FeO-NPs (green and chemical) followed by incubation for 24 hrs. To determine degraded effects of nanoparticles after incubation period, the samples were again quantified. It was estimated that nanoparticles have significant DNA degrading ability and DNA quantified as 154.9 ng/μl and 304.42 ng/μl respectively for green and chemically synthesized FeO-NPs.

### 3.7. Apoptosis, necrosis, and membrane permeability evaluation

Cell death was study through fluorescence microscopic analysis (AxioVert 200, Zeiss, Oberkochen, Germany) by using various fluorescent DNA binding dyes (ethidium bromide and acridine orange). Acridine orange (AO) permeates both viable and nonviable leishmanial cells, and emits green fluorescence by getting attached to the nucleic acid. Ethidium bromide (EB) permeates only nonviable cells, that have lost memebrane integrity, on intercalation into DNA, emits red fluorescence(Baskic, Popovic, Ristic, & Arsenijevic, 2006)

Fluorescent microscopic visualization showed that at earlier stage of incubation, about 80-90% FeO-NPs treated leishmanial cells radiating light green fluorescence showed early apoptosis. Cell emitting orange to red florescence, with fragmented chromatin material showed the late apoptosis. At the last stage of incubation, about 90% necrotic bodies emitting orange color were observed. (Fig.3.14).

**Fig 3.14.**
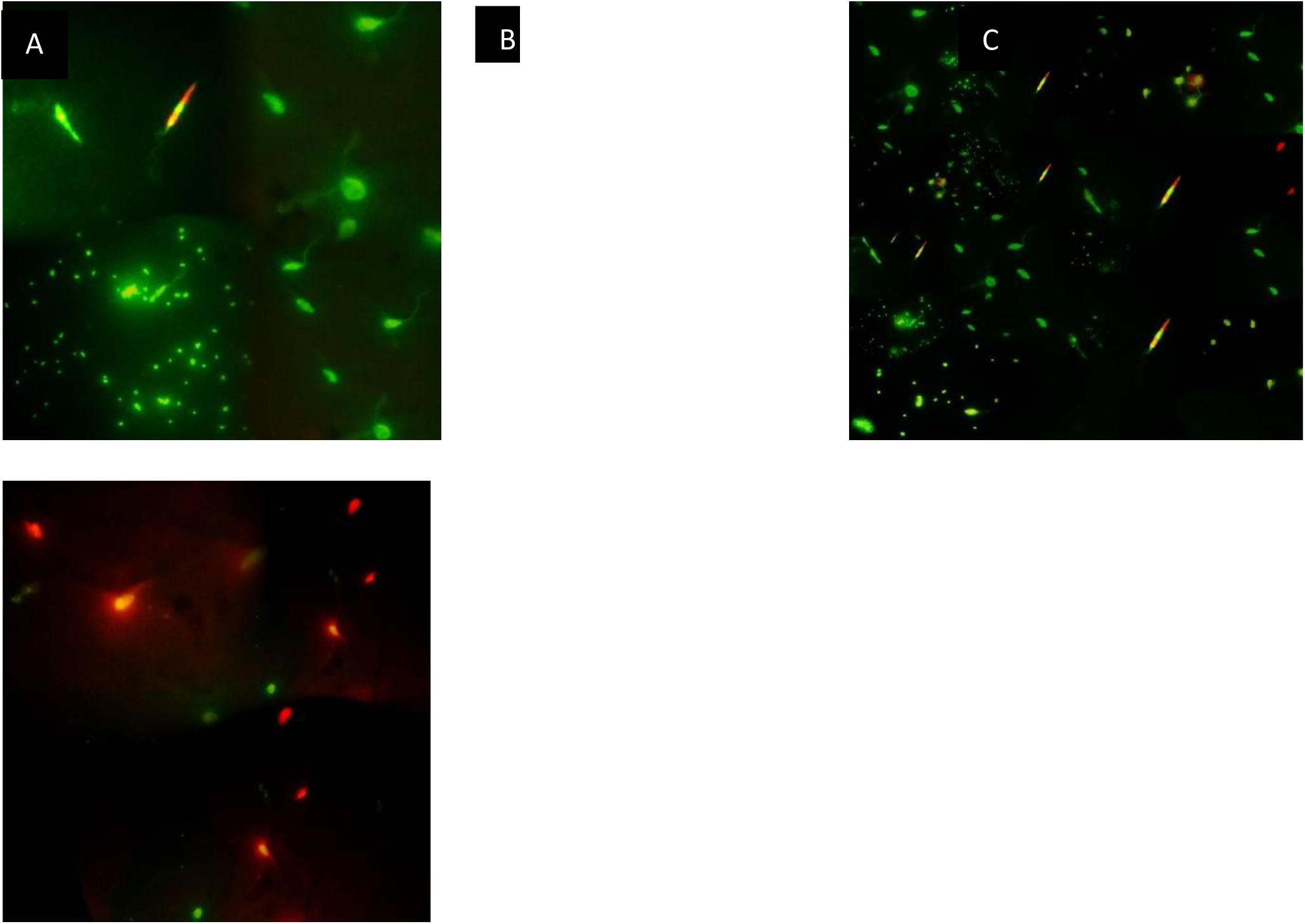
Apoptotic and necrotic bodies. Apoptotic and necrotic bodies of leishmanial cells after treatment with FeO-NPs for 24 hours. Notes: (A) Apoptotic bodies (B) Necrotic and apoptotic bodies (early and late apoptosis) (C) Necrotic bodies.

Membrane permeability evaluation was done by using SYTOX green dye. This dye has the ability to cross the cell membrane of compromised cells. Fluorescent microscopic visualization of FeO-NPs treated leishamanial cells emitting green color, confirmed the reactive oxygen species induced membrane integrity (Fig.3.15)

**Fig.3.15.**
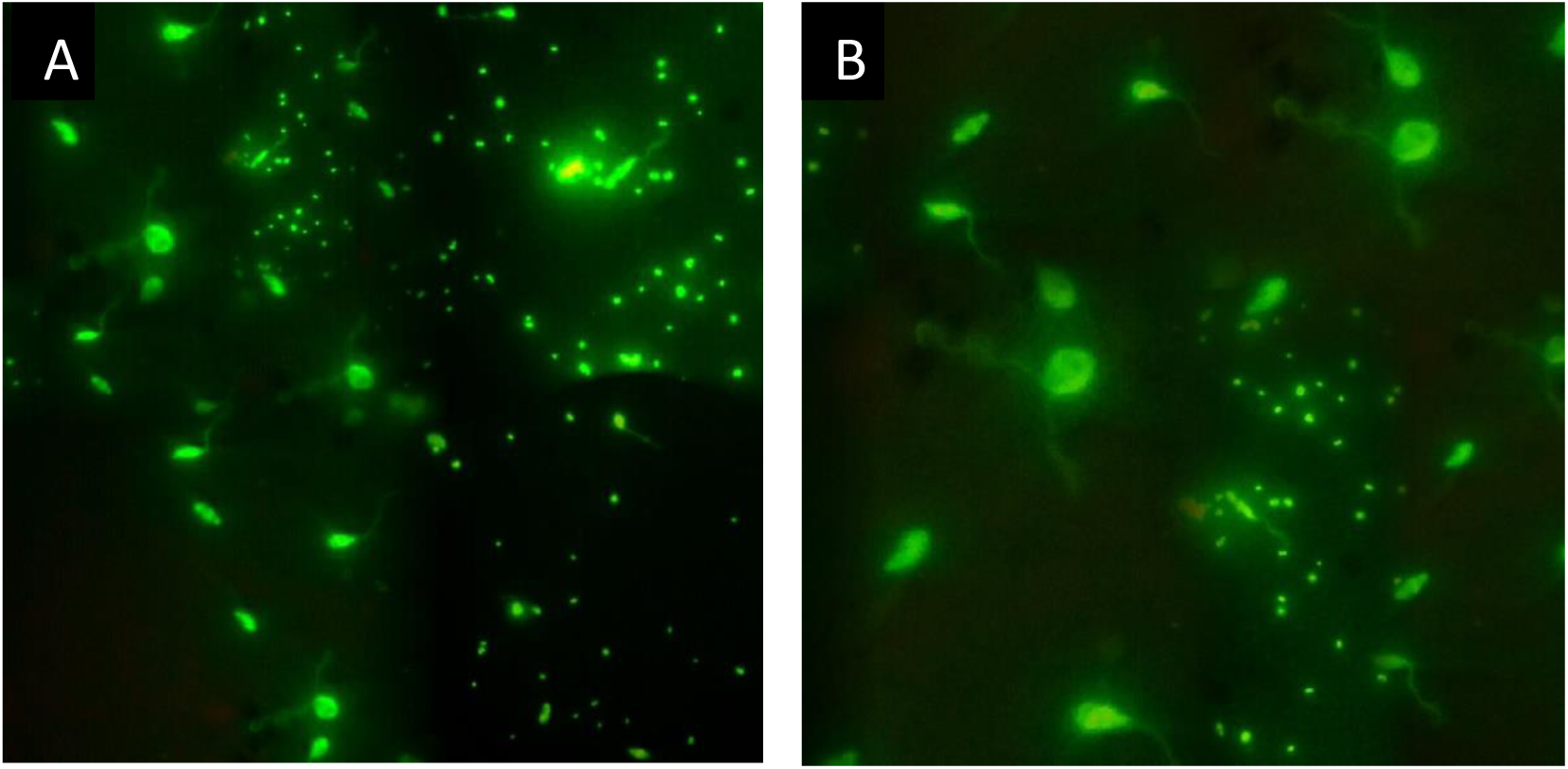
Fluorescent microscopy by using SYTOX green fluorescent dye (A) membrane permeability caused by Green synthesized FeO-NPs (B) By Chemically synthesized FeO-NPs.

## Conclusion

All the practice modalities for leishmaniasis lack appropriate safety, efficacy, and affordability. One of the most important issue associated with currently available leishmanicidals is the resistance by pathogens. There is need to develop more efficient candidates for fatal leishmaniasis. Green synthesis is best alternative remedy to synthesize eco-friendly metallic nanoparticles that avoids the use of toxic chemicals. Plant extracts contain certain important functional groups including citric acid, cyclic peptides, tannins, gallic acids and retinoic acid that not only causes bio-reduction but also act as capping and stabilizing agents. Plant mediated synthesis seems most reliable, facile, cost effective and eco-friendly route for metallic nanoparticles synthesis. Despite there being various studies regarding antibacterial, antifungal, antiviral efficacy of FeO-NPs have shown encouraging results. However, to date, in literature, there is no reported study regarding leishmanicidals effects of FeO-NPs. This study is the first to show the synergistic effects of FeO-NPs and LED light together demonstrating antileishmanial effects. The previous knowledge, regarding the sensitivity of *Leishmania* parasite against the ROS, encouraged to explore leishmanicidals properties of FeO-NPs for *L.tropica*, and comparative analysis of their efficacy under dark and light irradiation conditions. As it is estimated, LED irradiated, FeO-NPs exhibits significant antileishmanial properties against all promastigotes and amastigotes. It is demonstrated that this enhanced inhibitory effect under LED exposure, further DNA damage, membrane permeability, and cell death via apoptosis caused by FeO-NPs is directly related to generation of by reactive oxygen species by irradiating the nanoparticles. These ROS causes the condensation of chromatin material and nucleus shrinkage which led to the fragmentation of DNA, followed by the formation of apoptotic bodies eventually to the cell death.

## Notes

### Competing Interest Statement

The authors have declared no competing interest.

## References

Allahverdiyev, A. M., Abamor, E. S., Bagirova, M., Baydar, S. Y., Ates, S. C., Kaya, F., … Rafailovich, M. (2013). Investigation of antileishmanial activities of Tio 2@ Ag nanoparticles on biological properties of L. tropica and L. infantum parasites, in vitro. Experimental parasitology, 135 (1), 55–63.

Alvar, J., Velez, I. D., Bern, C., Herrero, M., Desjeux, P., Cano, J., … Team, W. L. C. (2012). Leishmaniasis worldwide and global estimates of its incidence. PloS one, 7 (5), e35671.

Alvar, J., Yactayo, S., & Bern, C. (2006). Leishmaniasis and poverty. Trends in parasitology, 22 (12), 552–557.

Baskić, D., Popović, S., Ristić, P., & Arsenijević, N. N. (2006). Analysis of cycloheximide-induced apoptosis in human leukocytes: fluorescence microscopy using annexin V/propidium iodide versus acridin orange/ethidium bromide. Cell biology international, 30 (11), 924–932.

Cornell, R. M., & Schwertmann, U. (2003). The iron oxides: structure, properties, reactions, occurrences and uses: John Wiley & Sons.

Fu, P. P., Xia, Q., Hwang, H.-M., Ray, P. C., & Yu, H. (2014). Mechanisms of nanotoxicity: generation of reactive oxygen species. Journal of food and drug analysis, 22 (1), 64–75.

Gleiter, H. (2009). Nanoscience and nanotechnology: the key to new studies in areas of science outside of nanoscience and nanotechnology. MRS bulletin, 34 (06), 456–464.

Mahdavi, M., Ahmad, M. B., Haron, M. J., Gharayebi, Y., Shameli, K., & Nadi, B. (2013). Fabrication and characterization of SiO 2/(3-aminopropyl) triethoxysilane-coated magnetite nanoparticles for lead (II) removal from aqueous solution. Journal of Inorganic and Organometallic Polymers and Materials, 23 (3), 599–607.

Matějka, V., & Tokarský, J. (2014). Photocatalytical nanocomposites: a review. Journal of nanoscience and nanotechnology, 14 (2), 1597–1616.

Nadhman, A., Nazir, S., Khan, M. I., Arooj, S., Bakhtiar, M., Shahnaz, G., & Yasinzai, M. (2014). PEGylated silver doped zinc oxide nanoparticles as novel photosensitizers for photodynamic therapy against Leishmania. Free Radical Biology and Medicine, 77, 230–238.

Naseem, T., & Farrukh, M. A. (2015). Antibacterial activity of green synthesis of iron nanoparticles using Lawsonia inermis and Gardenia jasminoides leaves extract. Journal of Chemistry, 2015.

Palmieri, B., & Sblendorio, V. (2007). Oxidative stress tests: overview on reliability and use. European review for medical and pharmacological sciences, 11 (6), 383–399.

Shah, N. A., Khan, M. R., & Nadhman, A. (2014). Antileishmanial, toxicity, and phytochemical evaluation of medicinal plants collected from Pakistan. BioMed research international, 2014.

Ullah, N., Nadhman, A., Siddiq, S., Mehwish, S., Islam, A., Jafri, L., & Hamayun, M. (2016). Plants as Antileishmanial Agents: Current Scenario. Phytotherapy Research.

Vankayala, R., Lin, C.-C., Kalluru, P., Chiang, C.-S., & Hwang, K. C. (2014). Gold nanoshells-mediated bimodal photodynamic and photothermal cancer treatment using ultra-low doses of near infra-red light. Biomaterials, 35 (21), 5527–5538.

Xiao, L., Gu, L., Howell, S. B., & Sailor, M. J. (2011). Porous silicon nanoparticle photosensitizers for singlet oxygen and their phototoxicity against cancer cells. ACS nano, 5 (5), 3651.

Yasinzai, M., Khan, M., Nadhman, A., & Shahnaz, G. (2013). Drug resistance in leishmaniasis: current drug-delivery systems and future perspectives. Future medicinal chemistry, 5 (15), 1877–1888.

Żamojć, K., Zdrowowicz, M., Rudnicki-Velasquez, P. B., Krzymiński, K., Zaborowski, B., Niedziałkowski, P., … Chmurzyński, L. (2017). The development of 1, 3-diphenylisobenzofuran as a highly selective probe for the detection and quantitative determination of hydrogen peroxide. Free radical research, 51 (1), 38–46.

